# An evolving cancer instigates clonally unrelated neighboring cells to form distant metastases

**DOI:** 10.1101/085423

**Authors:** Jie Dong, Cedric Darini, Farhia Kabeer, Sarah Dendievel, Junxia Min, Dolores Hambardzumyan, Louis Gaboury, Tom Lenaerts, Guillaume Darrasse-Jèze, Yi Li, Katrina Podsypanina

**Author notes:** These two authors contributed equally to this work (Jie Dong, Cedric Darini). Current address: Department of Molecular Oncology, British Columbia Cancer Agency, Vancouver, BC, Canada. Current address: Oncology Aflac Cancer and Blood Disorders Center, Emory University, Atlanta, GA, USA. Current address: UMR3644, Institut Curie, Paris, France (Katrina Podsypanina).

## Abstract

Based on the clonal evolution theory of cancer formation, a single cell within a tissue gains a cancer-driving mutation and thus a growth advantage. From this expanded cellular mass, another cell gains a new mutation allowing this newly mutated cell to gain new competitive advantage and to expand in number (thus clonal expansion). Another clone then emerges. Eventually all required mutations are gained, and a cancer forms. Consequently, while a primary lesion may harbor divergent subclones, all the subclones within the primary cancer as well as all metastatic growths in secondary organs share at least the very first oncogenic mutation that initiates the primary cancer. However, by tracking genetically marked mammary epithelial cells that suffered the initiating oncogenic mutation—and their neighboring mammary cells that did not-in several mouse models of human breast cancer, we found that genetically unrelated mammary epithelial cells can be colluded by neighboring mutated cells to disseminate, and that they can even undergo *de novo* tumorigenic transformation and form distant metastases. Therefore, clonally unrelated epithelial cells may contribute to cancer progression and to the heterogeneity of the systemic disease. The non-linear cancer spread has important implications in cancer prevention, treatment, and therapeutic resistance.

## Introduction

With the advance of large scale and single-cell sequencing, it is increasingly evident that cancer is heterogeneous and harbors multiple clonally evolving entities, which compete and sometimes collaborate in their access to nutrients, in resistance to therapy, and in colonization in distant organs. Systemic dissemination occurs usually after cells are fully transformed in the primary site but may also take place before all hallmarks of full malignancy are gained (Klein 2009; Rhim et al. 2012; Sanger et al. 2011; D. Weng et al. 2012). Premalignant cells are detected in the disseminated tumor cell pool in patients with cancer (Fiorelli et al. 2015; V. J. Hofman et al. 2011; V. Hofman et al. 2012). These disseminated precancerous/cancerous cells then may acquire new mutations conferring additional growth or survival advantages in distant organs (Gerlinger et al. 2012; Klein 2013; Melchor et al. 2014).

The premise of these studies and cancer clonal evolution theory in general is that all subclones within the primary cancer as well as all metastatic growths in secondary organs share at least the very first oncogenic mutation that initiates the primary cancer. However, histopathologically benign cells that are adjacent to an epithelial cancer also suffer genetic and epigenetic changes, and these cells can restrain or sometimes promote cancer evolution via paracrine signaling or direct cell-to-cell interactions (Eng et al. 2009; Hanahan and Coussens 2012). Direct recruitment of premalignant cells can also contribute to tumor progression at the primary site (Chapman et al. 2014; Cleary et al. 2014; Wu et al. 2010). Together, these observations suggest that cells that did not suffer the initial oncogenic mutation can participate in cancer formation and progression. Consistent with this possibility, some metastatic human tumors do not carry driver mutations that are present in the matched primary lesions (Kroigard et al. 2016; Mao et al. 2015). However, until now there is no direct experimental evidence that the benign neighbors contribute directly to metastatic cancer.

Breast cancer is the most frequently diagnosed cancer in women (except skin cancer) and the second leading cause of cancer death after lung cancer. Breast cancer arises from the epithelial cells of the breast ductal tree. We previously reported that normal mouse mammary cells injected into the tail vein of new hosts can colonize the lungs and, if they are later induced to express an oncogene, can form ectopic tumors (Podsypanina et al. 2008). Here, we demonstrated that fluorescently tagged normal mammary cells that were implanted orthotopically into mammary fat pads, consistently disseminated to the lung when disturbed by mammary tumors initiated by PyMT or ErbB2 transgene expression or by intraductal injection of 4T1 mammary cancer cell line. Next, we used intraductal injection of retrovirus to introduce either *Wnt1* or *ErbB2* into a small number of mammary epithelial cells to initiate mammary tumorigenesis, and found that some primary tumors contained an unrelated population of carcinoma cells that lacked the provirus DNA and the experimentally introduced oncogene. Importantly, we found that many of the metastatic foci in the lungs also lacked the provirus DNA and its oncogenic protein product.

## Results

### Normal mammary epithelial cells can be locally disseminated by adjacent mutated cells that are evolving into cancer

To establish whether the mammary cells that are clonally unrelated to primary tumor can be dispersed locally by their malignant neighbors, we studied chimeric mammary outgrowths formed by transplanting into the mammary fat-pad RFP-tagged normal mammary cells admixed with YFP-tagged mammary cells that also carried doxycycline-inducible PyMT (*ToMT:MTB*, Table S1). In the absence of doxycycline treatment (and thus no oncogene induction), a bi-color mammary tree formed after six weeks post-transplantation, and it was indistinguishable from the outgrowths produced by *RFP*-marked normal mammary cells that were admixed with oncogene-free, *GFP*-marked normal mammary cells (Fig S1A), confirming no leaky expression or abnormal behavior in these *ToMT:MTB:YFP* cells. After one week of doxycycline stimulation, these chimeric grafts developed numerous microscopic tumors (FFig 1A,B, S1B). Mitotic activity was detected in green PyMT-expressing cells, as expected; mitotic activity was also observed in the RFP+ PyMT-negative cells, though at a much lower frequency (Fig S1C-E). Furthermore, occasional solitary red cells were detected in the stroma (Fig 1B, S2A), which was reminiscent of cell delamination reported in early stages of cancer invasion (Rhim et al. 2012). Within more advanced tumors formed after 6 weeks of PyMT induction, RFP+ cell delamination was also frequently detected both by live fluorescence imaging and after immunostaining for RFP (Fig 1C,D S2B). Dispersion of normal cells marked by a different fluorophore (GFP) was similarly observed by confocal imaging of induced *ToMT:MTB:YFP/GFP* chimeric grafts (Fig S2C, Supplementary video S1). While transplanted tumor cells have been reported to affect branching of the endogenous mammary ductal tree (Parashurama et al. 2012), our data here showed local dissemination of the isolated benign mammary cells by their cancerous neighbors.

To investigate whether this local dissemination of the benign mammary cells may also occur during cancer formation in humans, we surveyed samples from patients with infiltrating lobular carcinoma (ILC). Normal breast epithelial cells produce E-cadherin, but the lobular carcinoma cells, beginning from the *in situ* stage, lack this marker because of genetic alterations in *CDH1*, which encodes E-cadherin (Berx et al. 1996), effectively making E-cadherin a biomarker of a benign state in ILC. The E-cadherin-positive and E-cadherin-negative cells have been reported in close association throughout different stages of disease (Andrade et al. 2012; Hwang et al. 2004; Mastracci et al. 2005; Vos et al. 1997). Extending these observations, we detected disrupted and infiltrating E-cadherin-positive cells among and E-cadherin-negative tumor cells in three out of five ILC cases (Fig 1E, Fig S3A,B). Interestingly, a recent study suggested that normal cell dissemination actually required E-cadherin (Shamir et al. 2014). Collectively, our findings in mice and humans documented architectural disruption of the benign epithelia that surround an evolving tumor.

**Figure 1.**
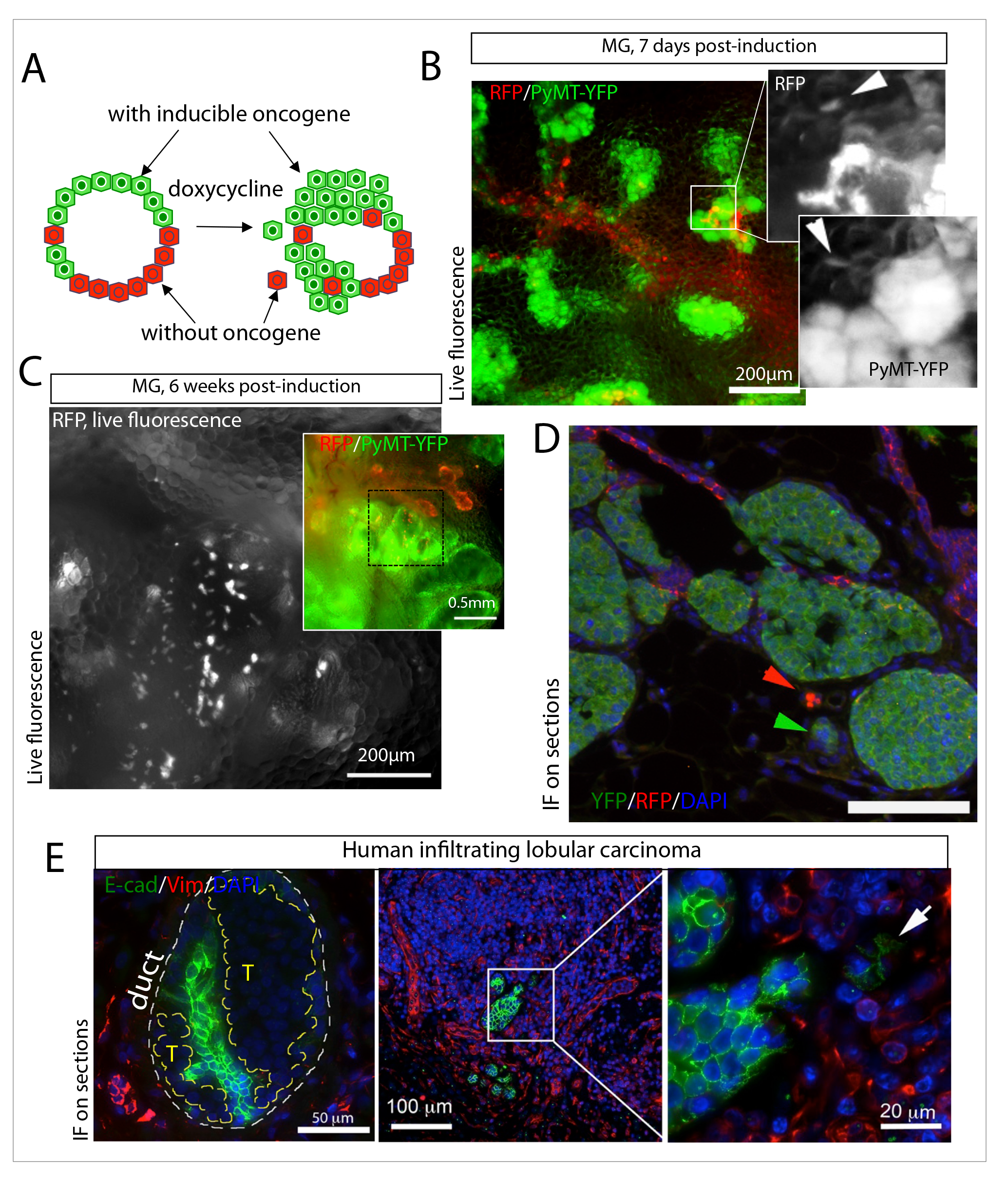
Local dispersal of normal mammary cells by growing breast tumors. **A.** A system for tracking parallel dissemination is based on mammary gland reconstitution from 2 genetically different donors. Schema illustrates tumor induction with doxycycline in cells of the donor carrying a doxycycline-inducible polyoma middle T oncogene and a YFP transgene (green), while cells from an oncogene-free donor, expressing RFP, remain untransformed (red). **B, C.** Normal mammary cells are scattered by growing tumorsin chimeric grafts. 5x10^6^ *ToMT:MTB:YFP* and 5x10^6^ RFP unsorted mammary cells were mixed and injected in equal doses in cleared inguinal fat pads of 5 *NOD.RagF^-/-^Il2rg^-/-^* (NRG) hosts. Hosts were placed on doxycycline for 7 days (**B**) or for 6 weeks (**C**) ten weeks after transplantation. Freshly dissected induced mammary glands were inspected under fluorescent microscope. Insets: B. Single-channel images of isolated cells spreading into fat pad (arrowheads); C. Composite color image of the field at lower magnification. MG-mammary gland. **D**. Scattering and delamination of normal epithelial cells in *ToMT:MTB:YFP/RFP* chimeric glands. Chimeric glands were prepared as described in (**B**). Chimeric mammary outgrowths were collected from mice exposed to doxycycline for 7 days, fixed, sectioned and stained with anti-RFP and anti-GFP antibodies. Red arrowhead points at normal cells outside of mammary duct, green arrowhead points at tumor cell cluster outside of mammary duct. IF – immunofluorescence. **E**. Delamination of normal epithelial cells in human lobular carcinoma samples. Left panel: E-cadherin+ cells (green) are delaminated from basal layer by E-cadherin-negative tumor mass (T) in human lobular carcinoma. White dashed line marks duct margins (duct); yellow dashed lines surround tumor cells (T) within the duct. Right panel: An isolated cluster of delaminated E-cadherin+ cells within an invasive lobular tumor mass. Inset: Arrowheads point at normal cells spreading into a tumor mass. E-cad = E-cadherin; Vim = vimentin.

### Under the influence of adjacent mutated cells that are evolving into cancer, neighboring mammary cells can disseminate to the lungs

To test whether benign mammary cells can be instigated by oncogene-activated cells to disseminate to distant organs, we transplanted mixtures of cells from *RFP* mice and from *ToMT:MTB:YFP* mice into the cleared fat-pads of immunodeficient mice, and examined the lungs of these recipients in the presence and in the absence of PyMT induction. As expected, without doxycycline treatment, no fluorescent cells were detected on the surface of the lungs in the three mice examined. In contrast, in the group treated with doxycycline for one week, RFP+ cells in singles or in aggregates were detected on lung surface in four out of five graft hosts by whole mount microscopy (Fig 2A). Using anti-RFP immunofluorescence staining of serial lung sections from five mice, seven positive cells and one positive cell cluster were detected (Fig S4A), while there were no lesions resembling metastatic nodules in the lungs at this early stage. Using a higher throughput technique, flow cytometry, about 0.29 ± 0.08% of size-gated cells were RFP+, and about 0.06 ± 0.01% were YFP+ in the doxycycline-induced cohort, while without doxycycline induction the proportion of red fluorescent cell remained within background limits (0.02 ± 0.02% red and 0.001 ± 0.001% green cells, Fig 2B, S5A). Moreover, the fluorescent cells were present within the EpCAM+ fraction, indicating that they were of epithelial origin (Fig 2C, S5B). We also found examples of E-cadherin+ disseminated RFP+ cells on sections (Fig 2D). The circulating fluorescent cells were detected at a lower frequency in the cardiac blood samples from the doxycycline-exposed animals (Fig. S5C,D), suggesting that the disseminated normal and tumor cells were more abundant than the circulating cells in this model. Importantly, the presence of ectopic cells of both colors after tumor induction suggested that tumor growth caused both PyMT-positive and PyMT-negative cells to disseminate. Thus, under the influence of an evolving mammary cancer, clonally unrelated and phenotypically benign mouse mammary epithelial cells disseminated to distant organs.

**Figure 2.**
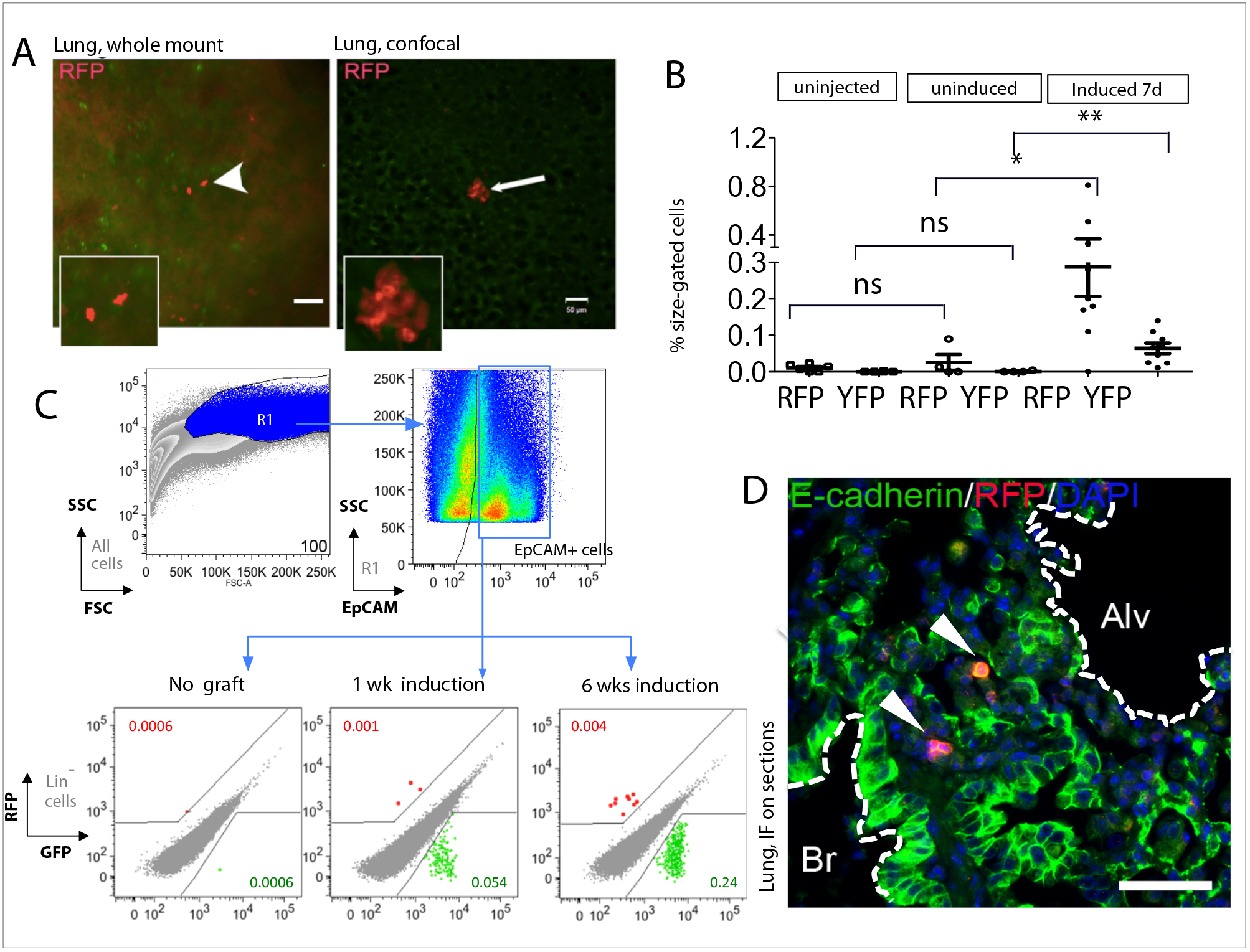
Systemic dissemination of normal mammary cells by mouse breast tumors. **A.** Isolated red cells (arrowhead) and a red cell cluster (arrow) observed in freshly harvested unperfused lungs from the chimeric gland hosts described in Fig. 1B. The faint green signals are detected uniformly throughout most fresh lung samples and likely represent autofluorescence. **C**. Dot plot showing percentages of YFP+ and RFP+ cells among size-gated cells in dissociated lung preparations from the chimeric gland hosts. Mammary cells from single *ToMT:MTB:YFP* and *RFP* donors were mixed as in **1B** and injected in equal doses in cleared inguinal fat pads of NRG hosts on 3 independent occasions. Hosts were placed on doxycycline 6-14 weeks after transplantation or left without doxycycline stimulation. Unperfused right lungs from mice exposed to doxycycline for 1 week (n=9), or left unexposed (n=4) and from graft-free controls (n=6) were prepared into single cell suspensions and evaluated by flow cytometry to measure the total number of YFP+ and RFP+ cells. Mann-Whitney test *p*-value<0.001(**) and <0.01(*). **C.** Gating strategy and representative flow cytometry scatter plots showing that ectopic cells in the lungs of doxycycline-stimulated chimeric gland hosts express epithelial cell marker EpCAM. Unperfused right lungs from mice exposed to doxycycline for 0 (n=5), 7 days (n=3) or 6 weeks (n=3) were prepared into single cell suspensions and stained with anti-CD3, -CD19, -CD11b, -F4/80 (Lin), anti-Ter119, anti-EpCAM and anti-CD29 antibodies. Lung preparations were then evaluated by flow cytometry. Lineage exclusion was applied to avoid detection of fluorescent hematopoietic cells resulting from carryover of white blood cells from the donor MG preparation into immunodeficient hosts. The percentages of fluorescent cells among EpCAM+ cells are shown in red and green. **D**. RFP-E-cadherin double positive cells in a lung section from formalin-fixed, paraffin embedded (FFPE) tissue described in **2A**. Arrowheads point at the RFP+ cells. White line highlights bronchial lumen (Br) and alveolar space (Alv).

To determine whether this distant dissemination of normal mammary cells from mammary glands with an evolving cancer also occurred when the instigating tumors were initiated by an oncogene commonly found in human breast cancer patients, we transplanted benign mammary cells expressing GFP admixed with mammary cells with inducible expression of activated Her2/Neu/ErbB2 (*TAN:MTB* mice), a potent human oncogene which is amplified and overexpressed in 20-25% of human breast cancer and encodes a member of the epidermal growth factor receptor family of receptor tyrosine kinases (Slamon et al. 1987). Again, GFP-positive cells were detected in the lungs of doxycycline-stimulated syngeneic recipients, showing linear relationship between the proportion of GFP+ cells in the chimeric mammary glands and in the lungs of individual animals (Fig S5E,F). Collectively, these *in vivo* data using two genetic models suggested that during breast cancer progression, the disseminated cell pool included both cancerous mammary epithelial cells with the initiating oncogene and phenotypically benign mammary cells without the initiating oncogene.

### The dynamics of disseminated mammary cells in the lung

We next examined the dynamics of benign cell dissemination in the 4T1 intraductal transplant model. First, we intraductally injected into normal mice RFP+ normal mammary cells and the 4T1 mouse mammary cancer cells, separately (n=9) or admixed (n=12). Two weeks later mammary tumors formed in mice injected with 4T1 cells (Fig 3A), and the RFP+ benign cells co-transplanted with 4T1 cells became highly disorganized and scattered within the massively expanded tumors (Fig 3B). In contrast, in glands injected with RFP+ cells alone, RFP+ cells lined up and integrated in the intact mammary ducts (Fig S6A). Nevertheless, the RFP+ cells with or without tumors retained their epithelial nature based on positive staining for E-cadherin (Fig S6B,C). By flow cytometry, similar proportions of RFP+ cells were detected in mammary tissues of mice with and without tumors, while no RFP+ were detected in the animals injected with 4T1 tumor cells only (Fig 3C), indicating that RFP+ cells persisted in the mammary gland regardless of co-existing tumor cells.

Next, we excised chimeric tumors in four of the eight remaining animals. After another two weeks, we examined the lungs of all eight mice. Pulmonary metastases were detected in all mice, suggesting that tumor cells had disseminated prior to the removal of the primary tumor (Fig S6D). Importantly, flow cytometry readily detected RFP+ benign mammary cells in the lungs of the mice that received both RFP+ cells and 4T1 cells, but not in the lungs of the control cohorts of mice (Fig 3D,E), in agreement with our findings in mice with doxycycline-induced tumors described above. Staining with E-cadherin confirmed the epithelial origin of the RFP+ cells sorted from the primary tumors and lungs of these mice (Fig S6E).

**Figure 3.**
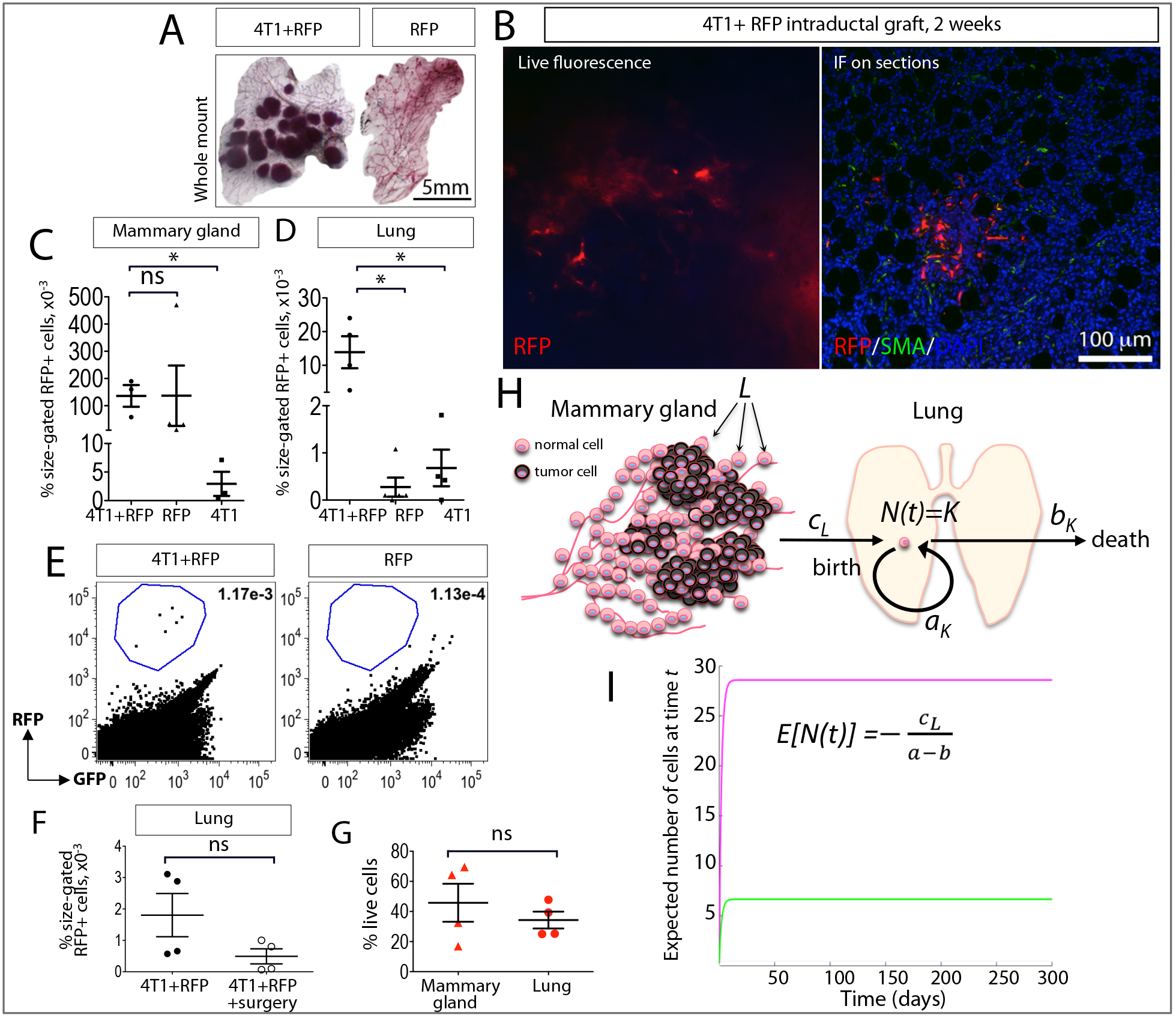
Mammary cell dynamics in the lung. **A.** Intraductal injection of 4T1 cells gives rise to mammary tumors. 5xl0^4^ 4T1 cells and/or 5x10^4^ *RFP* unsorted mammary cells were mixed and injected in the thoracic and inguinal mammary ducts of 12 *NOD.ScidJI2rg^−/−^*(NSG) hosts on two independent occasions. Animals injected with pure 4T1 or pure RFP cells served as control (n=9). Animals were followed for two (left) or for four (right) weeks. Freshly dissected mammary glands were fixed and stained with carmine to detect mammary ducts and tumors. **B**. Local scattering of normal donor mammary cells by 4T1 tumor cells. Tumor-bearing mammary glands were collected at two weeks and examined for RFP fluorescence (left panel), then fixed, sectioned and stained with anti-RFP and anti-SMA antibodies (right panel). IF – immunofluorescence. **C**. Intraductally injected RFP+ cells persist in the tumor-bearing and tumor-free mammary glands. Mammary glands from mice injected with 4T1, RFP, or a mixture of 4T1 and RFP cells as described in 3A were prepared into single cell suspensions four weeks after injection and evaluated by flow cytometry. Dot plot shows percentages of RFP^+^ cells among size-gated cells in dissociated mammary preparations. LTnpaired t test *p*-value>0.01 (ns), <0.01(*). **D**, **E**. Normal RFP cells disseminate to the lungs only in the presence of a growing tumor. **D**. Dot plot showing percentages of RFP^+^ cells among size-gated cells in dissociated lung preparations from mice injected with 4T1 (n=4), RFP (n=5), or a mixture of 4T1 and RFP cells (n=4). Mann-Whitney test *p*-value<0.01(*). **E**. Representative flow cytometry scatter plots from mice described in 3D. The percentages of red cells are shown in the upper left corner. **F**. Spontaneously disseminated normal cells do not accumulate in the lungs. Bar plot showing percentages of RFP+ cells in dissociated lung preparations from animals evaluated two weeks after surgery, four weeks after injection, and from untreated controls evaluated before surgery, two weeks after the injection. Mann-Whitney test *p*-value>0.01(ns). **G**. Viability of RFP+ cells in cell suspensions from the mammary tissue and from the lung. Mann-Whitney test *p*-value>0.01(ns). **H**. Schematic representation of the birth-death process model applied to breast cells dissemination to the lungs. *L*, number of normal-tumor cell contacts in the breast; *c_L_*, dissemination rate from the breast when there are *L* contacts in the breast; *N(t)*, number of marked cells at time *t* in the lungs; *K* is an integer value; *a_K_*, birth rate when there are *K* cells in the lung; *b_K_*, death rate when there are K cells in the lung. *a_K_*+*c_L_*= *λ_K_*, which is the combined birth rate used in the Methods section. **I**. Growth curve simulation shows that the mean number of disseminated cells becomes a multiple of *c_L_* within a short time from the start of migration. Parameter values: *a* = 0, as most disseminated cells do not replicate in the lungs, *b* = l/2, as most disseminated cells disappear within 2 days of arriving in the lung; *c_L_*: magenta = 14.3, meaning 100 cells per week; green = 3.33, meaning 100 cells per month.

Interestingly, while in the mice with un-resected tumors the proportion of disseminated benign RFP+ cells among size-gated cells had reached 13.9±4.8 per 1×10^5^ by four weeks post-injection (Fig 3D), the proportion of RFP+ cells in the lungs of mice with surgically resected tumors had slightly decreased in comparison with the values measured at two weeks post-injection (1.8±0.7 per 1×10^5^ vs. 0.5±0.2 per 1×10^5^, respectively, Fig 3F). This finding suggests that the benign mammary cells continuously disseminated from the primary site, but did not proliferate after reaching the lungs over this two-week window of time. We also noted that the proportion of viable cells, as determined by DAPI exclusion, was modestly, but not significantly, lower among the disseminated RFP+ cells in the lungs than among RFP+ cells persisting in the primary tumors (34.3±5.6% vs. 45.8±12.6%, respectively, Fig 3G). Thus, many of the spontaneously disseminated normal mammary cells either failed to proliferate or died during the post-surgery follow-up period.

Based on this behavior of mammary epithelial cells at an ectopic site, we built a mathematical model to describe the dynamics of the disseminated cells in distant organs (Fig 3H,I). The presence of ectopic cells in the lungs was translated into a birth-death equation, describing temporal changes in cell numbers as a function of cell division, cell migration and cell death, as described in Materials and Methods. To estimate the number of mammary cells that may be in the lungs at a given time point, we simulated ectopic cell growth curves for dissemination rates of 100 cells per week and per month (Fig 3I, magenta and green lines, respectively). This simulation confirmed that from the start of constant cell efflux from the mammary gland, the ectopic cell count in the lung reaches an equilibrium within a short period of time irrespective of the dissemination rate. More broadly, this mathematical model suggests that normal cells detected in circulation of patients with cancer (Fiorelli et al. 2015; V. J. Hofman et al. 2011; V. Hofman et al. 2012) constitute evidence of sustained non-linear dissemination from the human tumors.

### Under the influence of adjacent mutated cells that are evolving into cancer, mammary epithelial cells without the initiating oncogene can form distant metastases

Having established that clonally unrelated mammary cells can be instigated by an evolving cancer to continuously disseminate to distant organs, we next asked whether under the influence of an evolving cancer, mammary epithelial cells without the initiating oncogene can not only spread, but also grow and progress into metastatic tumors. We previously used intraductal injections of the RCAS viral vector to deliver oncogenes into the luminal cells in mice transgenic for *tva*, which encodes the cytoplasmic membrane receptor for RCAS (Du et al. 2006). Proviral integration into the genome of mammary cells allows us to differentiate the transduced, oncogene-expressing cells from the neighboring non-transduced cells. Intraductal injection of RCAS expressing a constitutively activated, HA-tagged ErbB2 (RCAS-*caErbB2*) into virgin mice that expressed the *tva* transgene from the mammary epithelium-selective MMTV promoter leads to tumor development with a median latency of three months (Haricharan et al. 2014). We examined H&E-serial sections of the lungs from 31 tumor-bearing mice and detected 20 metastases in 13 mice. Using an antibody to detect the HA tag in the *caErbB2* oncoprotein, we found that while 13 metastatic lesions contained the HA tag in all tumor cells, seven metastases from six different mice were negative for the HA tag (Fig 4A, S7A,C). Moreover, genomic DNA from lung sections with HA-negative foci did not contain the caErbB2 oncogene or the provirus, while a strong signal was detected by qPCR in DNA from lung sections with HA-positive foci (Fig 4B). Therefore, most or all of the cells in HA-negative lung metastases were descendants of cells that were never infected by RCAS-*caErbB2.*

To assess metastatic heterogeneity in genetically more complex tumors, we examined metastatic mammary tumors in mice lacking *p19Arf*, a tumor suppressor gene which is commonly silenced in breast cancer (Silva et al. 2003). *caErbB2* overexpression cooperates with *Arf* loss, resulting in accelerated tumor development in *Arf*-null mice infected with RCAS-*caErbB2* (Sinha et al. 2015). Lung metastases were detected in six out of 13 Arf-null mice. Among 10 metastatic lesions from the *Arf*-null mice examined, five were negative for *caErbB2*-HA (Fig S7B), validating our observations in the *caErbB2*-only model.

We next examined the primary tumors for the contribution of the mammary cells that did not gain the initiating ErbB2 oncogene. Among 19 primary tumors examined (from both *Arf*-null and wild-type genotypes), the HA-tag was detected in all epithelial cells in nine tumors, including the three tumors from animals with HA-negative lung metastases (Fig 4C,D). A heterogeneous HA expression pattern was detected in the remaining 10 cases (Fig 4C,D), suggesting that ErbB2-initated tumorigenesis involved mammary epithelial cells that did not gain the initiating ErbB2 oncogene.

Previously, we found that oncogenic activation of ErbB2 causes aberrant proliferation and DNA damage (Reddy et al. 2010), which is known to lead to mutational activation of tumorigenic pathways. To investigate whether mammary cells in the vicinity of RCAS-*caErbB2*-uninfected cells may also experience DNA damage, we immunostained sections from RCAS-caErbB2-infected glands for γH2A.X, a common marker of DNA damage. Both HA-negative and HA-positive epithelial cells in RCAS-*caErbB2*-induced early lesions displayed an increased proportion of γH2A.X-positive nuclei over that observed in the uninjected control mammary glands (Fig 4E,F, S8A), suggesting an ongoing DNA damage in both infected and in the neighboring non-infected cells. However, we did not find any γH2A.X-positive nuclei in the RFP+ benign mammary cells scattered within tumors produced by inducible PyMT expression, or in RFP+ benign cells neighboring the 4T1 tumors (Fig S8B-D). Thus, while the benign mammary tumor neighbors disseminated to the lungs of tumor-bearing mice in all three models, their susceptibility to DNA damage and subsequent mutations depended on experimental context and the primary tumor type.

**Figure 4.**
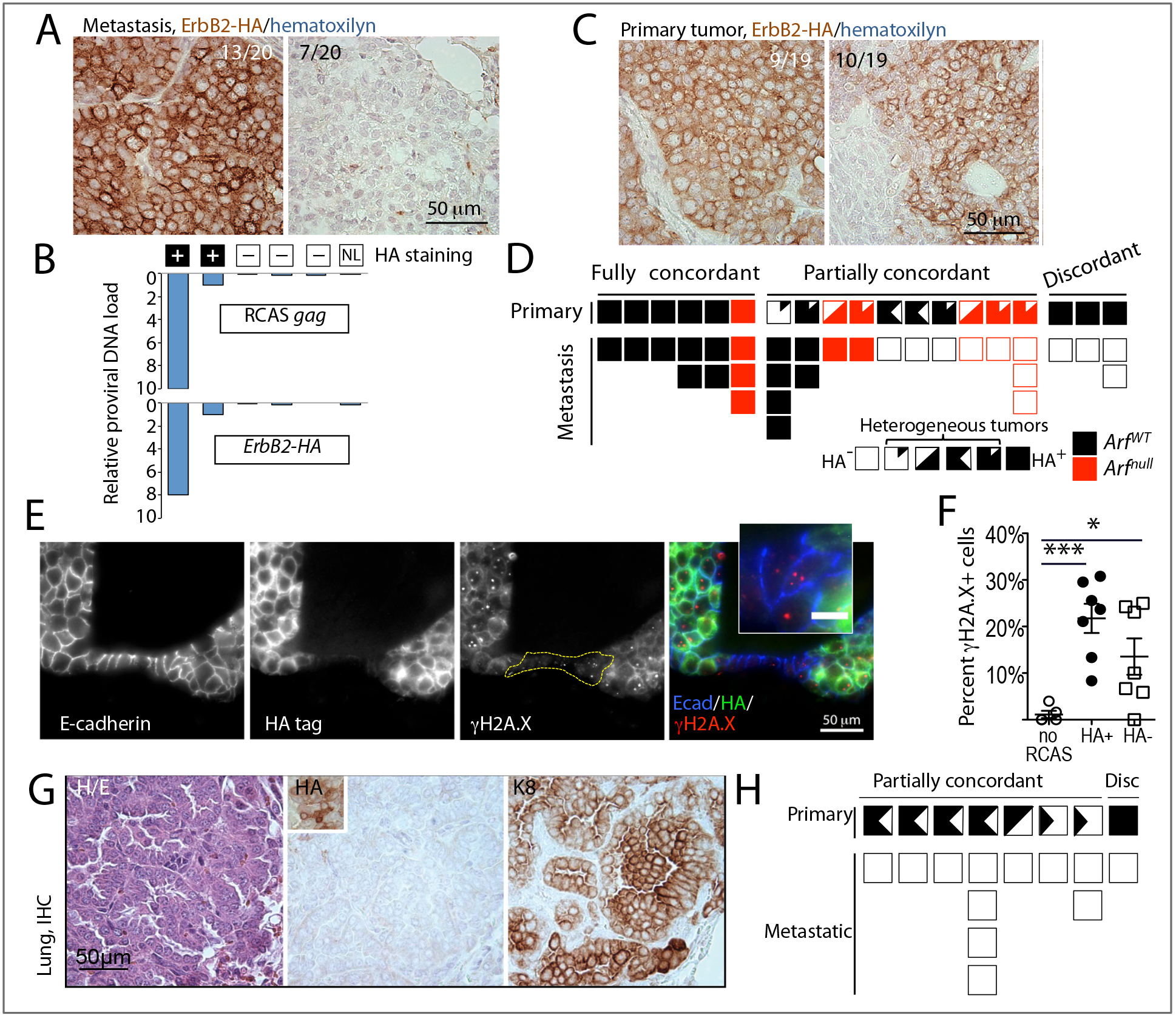
Metastatic tumors are initiated by mammary cells unrelated to tumor cells infected with oncogene-containing RCAS virus. **A.** Individual metastases in mice with RCAS-caErbB2-induced mammary tumors lack ErbB2-HA expression. Representative images of spontaneous lung metastases stained with anti-HA antibody. Left panel, HA+ (brown stain); right panel, HA-cells. MG-mammary gland, IHC-immunohistochemistry. **B**. Absence of proviral DNA in the HA-lung metastasis samples. Proviral load was calculated based on qPCR results using RCAS *gag*-specific (top graph) and *ErbB2/HA*-specific (bottom graph) oligos on DNA extracted from metastasis-containing lung sections. Lung sections containing HA+ lung metastases served as positive controls, normal lung (NL) from an uninfected mouse served as a negative control. **C**. Heterogeneity of *ErbB2*-HA expression in a fraction of RCAS-*caErbB2*-induced mammary tumors. Representative images of primary tumors with uniform (left panel) and patchy (right panel) pattern of anti-HA staining. **D**. Discordant ErbB2-HA expression in primary-metastatic tumor pairs from *Arf^WT^*(black boxes) and *Arf^null^* (red boxes) mice bearing RCAS-*caErbB2*-induced mammary tumors. Filled boxes-HA+, clear boxes – HA-. Heterogeneous cases are represented by partially filled/clear boxes. **E**. γH2A.X foci (red) in mammary cells in HA+ (green) and HA-(not-green) epithelial mammary cells (E-cadherin+, blue). **F**. Scatter plot showing percentages of γH2A.X+ cells in HA+ and HA-from early RCAS-ErbB2-induced lesions. Early lesions containing adjacent HA+ and HA-cells (n = 7) or normal ducts from uninjected glands (n = 5) from three mice were sampled. Unpaired t-test *p*-value<0.0001(***) and <0.01(*). **G**. Individual metastases in mice with RCAS-Wnt/-induced mammary tumors lack *Wnt1*-HA expression. Representative images of consecutive sections from a spontaneous metastasis stained with: left panel, hematoxylin/eosin (H/E), center panel, anti-HA antibody (inset shows HA+ cells in the paired primary tumor); right panel, anti-keratin 8 (K8) antibody. **H**. Discordant *Wnt1*-HA expression in primary-metastatic tumor pairs from mice bearing RCAS-*Wnt1*-induced mammary tumors. Filled boxes – HA+, clear boxes – HA-. Heterogeneous cases are represented by partially filled/clear boxes.

To validate the above observation of non-linear cancer progression in another model of breast cancer, we examined pulmonary metastases from mice bearing primary tumors initiated by RCAS carrying the gene encoding Wnt1, which activates a pathway commonly dysregulated in triplenegative or basal-like breast cancer and has been reported to induce mammary tumors with pulmonary spread in mice (Bu et al. 2013; Li et al. 2000). We injected RCAS expressing HA-tagged *Wnt1* (RCAS-*Wnt1* (Dunn et al. 2001)) intraductally into 55 mice that expressed the *tva* transgene from the mammary epithelium-selective MMTV or WAP promoter. Eleven mammary tumors developed with a median latency of 10 months, and *Wnt1*-HA+ cells were detectable in both models (Fig S9A). However, ten out of 11 RCAS-Wnt1 tumors also contained variable proportions of HA-negative tumor cells (Fig S9B,C, left panel), while only one tumor was homogeneously HA-positive (Fig S9C, right panel). We detected keratin 8-positive metastases in lung sections from eight out of these 11 mice (Fig 4G). All 12 metastatic tumors from these eight mice were negative for the HA tag, including the one case from the mouse bearing the primary tumor with homogeneous HA+ staining (Fig 4H). We also did not detect the oncogene or the RCAS provirus in genomic DNA from any of the metastasis-containing sections, irrespective of whether they originated in the MMTV-*tva* or the WAP-*tva* transgenic line (Fig S9D). Therefore, these lung metastases descended from mammary epithelial cells that were not infected by RCAS-*Wnt1*. Collectively, our observations in three distinct RCAS models strongly indicate that under the influence of mammary epithelial cells suffering oncogenic mutations and undergoing tumorigenesis, the neighboring normal mammary epithelial cells without the initiating oncogene can also evolve into cancer cells-likely by suffering DNA damage and gaining new oncogenic mutations-and can even spread to distant organs and to progress to frank metastases.

## Discussion

Using several mouse models, we demonstrated that an evolving cancer can provoke local and systemic dissemination of its epithelial neighbors, culminating in their clonal expansion not only in the primary site but also in distant locations. These unexpected observations suggest that mammary cells prior to the gain of an oncogenic driver can be instigated by neighboring cancerous cells to disseminate, and that epithelial cells that are clonally unrelated to an evolving cancer not only can contribute to the primary tumor, but also can become the principal source of distant metastases. Thus, tumor progression may be a non-linear process involving epithelial cells with different transformation histories.

Using two different transplantation techniques, we engineered chimeric models comprised of genetically marked normal and tumor cells, and showed that benign epithelial cells can participate in a growing tumor and can even disseminate to the lungs along with tumor cells. These findings are in agreement with the abundant clinical evidence that noncancerous epithelial cells can disseminate, for example, following surgery (Crisan et al. 2000; Seiden et al. 1994) and mechanical stimulation (Diaz et al. 2004), and during inflammation (Murray et al. 2013; Pantel et al. 2012). Premalignant cell dissemination has long been suspected to occur in patients with malignant tumors—as the peripheral blood of some cancer patients harbors epithelial cells with benign phenotypes (Fiorelli et al. 2015; V. J. Hofman et al. 2011; V. Hofman et al. 2012). Our mathematical simulations argue that the presence of precancerous cells in circulation in patients with established tumors may constitute evidence of sustained non-linear dissemination in human cancer. It has been proposed that cells that disseminate at the “neoplastically immature” state may undergo independent neoplastic progression after colonization, potentially complicating the course of the disease and the choice of therapies (Klein 2013).

Oncogene-activated cells may promote dissemination of clonally unrelated cells by several potential mechanisms, including mechanical pressure (Fernandez-Sanchez et al. 2015), collective invasion (Carey et al. 2013; Cheung et al. 2013), or by transmission of dissemination-promoting factors via extracellular vesicles (Zomer et al. 2015). Inter-clonal collaboration has been reported between Ras-mutated cells and scribbled-null cells where JNK signaling-mediated upregulation and release of certain cytokines in one cell activates JAK-STAT signaling in another cell (Wu et al. 2010). Therefore, as a downstream component of ErbB2 or PyMT signaling, JNK may cause oncogene-expressing cells to release cytokines to stimulate JAK-STAT signaling in neighboring benign epithelial cells. JAK-STAT signaling is known to stimulate breast cancer risk and metastasis (Haricharan and Li 2014). On the other hand, Wnt signaling has long been recognized as a key mediator in cell-cell interactions during cancer evolution (Cleary et al. 2014; Thliveris et al. 2013). Wnt ligands and other factors secreted by RCAS-Wnt1-infected mammary epithelial cells could cause genetic and epigenetic changes in non-infected mammary cells that promote dissemination.

The clonal expansion of the benign cells colluded by cancerous neighbors is likely driven by new genetic and epigenetic changes that are acquired before or after dissemination. In this study, we found evidence that introduction of an oncogene into mammary glands with an avian retroviral vector initiates and/or promotes DNA damage in both the infected and in the non-infected mammary cells. These genomic damages may lead to gains of oncogenic mutations in the benign cells that promote survival in distant sites. We did not detect evidence of DNA damage in the benign epithelial cells present in transgene-initiated or cell-line-derived tumors, perhaps explaining the absence of significant contribution of these naïve cells in metastatic progression of these tumor models.

Cancer metastasis is usually a highly inefficient process. Among fully transformed cells, only a small fraction goes on to form distant metastases. For example, in transgenic mice, a substantial proportion (20-30%) of animals remains metastasis-free even after having developed multiple large tumors driven by potent oncogenes, such as activated HER2/Neu (Muller et al. 1988) or PyMT (Guy et al. 1992)). The vast majority of cancer cells entering the metastatic process fails to form ectopic tumors, manifesting “metastatic inefficiency” (Mehlen and Puisieux 2006; Weiss 1990). It is likely that distant colonization requires factors and pathways that are distinct from the ones provided by the oncogenic drivers that cause primary cancer, and that most established malignant cells may lack the traits that are essential in distant colonization. Our observations suggest that a culprit to distant metastasis may sometimes lay in a clonally unrelated cellular compartment.

In both RCAS-ErbB2 and RCAS-Wnt1 models, benign mammary cells that did not apparently suffer an oncogene mutation unexpectedly showed equal or elevated proficiency in progressing into metastases compared to their neighbors that were forced to express an oncogene. One possible explanation is that the oncogenic drivers in these models regulate certain factors that hinder dissemination, for example, by promoting a more luminal-restricted cell fate (Lawson et al. 2015; Li et al. 2003; Vaillant et al. 2008). Alternatively, these oncogene-activated cells may be less apt to persist in the lung environment due to elevated anticancer barriers (apoptosis and senescence), but these barriers remain inactive in the mammary cells that do not express these potent oncogenes (Reddy et al. 2010). Human studies show that histologically benign, tumor-adjacent epithelia may nevertheless carry independent somatic mutations, including those in known cancer genes (Cooper et al. 2015; Forsberg et al. 2015; Z. Weng et al. 2015). These benign cells may also disseminate and contribute to distant metastases. This possibility explains why some metastases lack apparent cancer drivers found in the primary site. For example, virtually all studies evaluating HER2 discordance in primary vs. metastases detected HER2 FISH-negative metastases in patients with a HER2 FISH-positive primary tumor (Arslan et al. 2011; Houssami et al. 2011; Niikura et al. 2012; Santinelli et al. 2008; Simon et al. 2001; Vignot et al. 2012). Discordance was detected in patients both prior to and after the widespread use of anti-HER2 therapy (Vincent-Salomon et al. 2002; Xiao et al. 2011). Our findings in RCAS-caErbB2-HA-infected mice argue that the mismatched ERBB2/HER2 metastases in the clinic may be initiated by breast cancer neighbors that have never amplified HER2 in the first place.

In conclusion, our work in mice shows that mammary tumorigenesis can cause clonally unrelated epithelial neighbors to participate in cancer development both at the primary site and at distant locations. The evidence for a non-linear evolution of cancer and for a non-linear origin of cancer metastases in several models of human breast cancer offers a new interpretation of primary and metastatic tumor heterogeneity, and suggests that the “adjacent normal” tissue is a reservoir for cancer progression and spread and that it perhaps should also be targeted in cancer prevention and treatment.

## Materials and methods

### Animal Husbandry and Genotyping

All animal lines, listed in Table S1, were kept in specific pathogen-free housing with abundant food and water under guidelines approved by the Institut de recherches cliniques de Montréal Animal Care and Use Committee in accordance with the regulations of the Canadian Council on Animal Care, or by the Institutional Animal Care and Use Committee of Baylor College of Medicine, Houston, TX, USA. Doxycycline was administered by feeding mice with doxycycline-impregnated food pellets (625 ppm; S5829, Bio-Serv). Mice were placed on doxycycline as indicated for specific experiments. Animals were inspected for health weekly according to institutional guidelines.

### Mammary cell dissociation

Mammary glands were dissected from 6 to 16-week-old virgin female mice, minced with scissors and incubated with Collagenase/Hyaluronidase mix (StemCell Technologies) for two hours with trituration; or dissected and incubated with 3 mg/ml collagenase (Roche Diagnostics) overnight in a shacking 37°C incubator. Collagenase-digested samples were washed and strained through a 40μm filter, then treated with trypsin (0.25% Lonza) for 1-4 min. Then cells were washed with ten volumes of 0.1% w/v soybean trypsin inhibitor (Sigma #T9128) in PBS to stop the reaction, resuspended in PBS, counted and used for injections.

### Mammary cell culture

Freshly prepared mammary cell suspension were plated in 6 well plates and placed at 37°C in a CO_2_ incubator with 1.5 mL of supplemented serum-free medium (Mammary Epithelial Cell Medium BulletKit, containing one 500-mL bottle of Mammary Epithelial Cell Basal Medium and supplements: 2 mL of bovine pituitary extract, 0.5 mL of hEGF, 0.5 mL of hydrocortisone, 0.5 mL of GA-1000, 0.5 mL Insulin (Lonza)). Next day, un-adhered cells were transferred into a new plate and fresh media was added to the adherent primary mammary culture. On the 2^nd^ day, all un-adhered cells were discarded, and adherent fraction was detached with 0.1 mL 0.05% Trypsin-EDTA (Sigma). Then cells were washed with 3 mL 0.1% w/v soybean trypsin inhibitor (Sigma) in PBS to stop the reaction, and then resuspended in PBS, counted and used for injections.

### 4T1 cell culture

Frozen 4T1 cells were defrosted and cultured in T75 flasks at 37°C in a CO_2_ incubator with 10 mL of DMEM/F12 media with L-glutamine supplemented with 10% FBS. Cell were detached with Versene, washed, resuspended in PBS and counted.

### Fat pad injections

Mammary glands were surgically cleared of the endogenous mammary epithelium as described (Smith 1996) and injected with donor cells as indicated for specific experiments.

### Intraductal injections

Mammary cells were delivered intraductally as described (Du et al. 2006).

### Live fluorescence microscopy

Freshly dissected tissues were placed between glass slides and inspected under the fluorescent microscope (Nikon, Leica) or the confocal inverted microscope LSM510 (Zeiss). Then grafts were fixed in 10% buffered formalin for 16-48 h at room temperature, placed in 70% ethanol, and sent for paraffin embedding and sectioning at the histology platform at the IRCM, or at EnvA, France, or dissociated as described above.

### Virus preparation and delivery

RCAS virus was prepared and delivered intraductally as described (Du et al. 2006).

### Lung harvest and flow cytometry

In mice with chimeric grafts, freshly dissected, un-perfused whole lungs were inspected under fluorescent microscope as described above. Left lungs were fixed and sent to be sectioned as described above, and right lungs were minced with scissors and enzymatically digested as for mammary gland dissociation. Mammary cells from RFP, GFP or FVB mice were collected as control for surface profiling experiments. Single cells were stained with antibodies listed in Table S2 for 20 min on ice, washed, and resuspended in PBS:2%FBS before analysis. Flow cytometry was performed using an LSR or Fortessa (BD Biosciences) equipment at the IRCM or hôpital Necker platform. In mice with RCAS-induced tumors, lungs were fixed in 4% paraformaldehyde overnight at 4 °C, paraffin-embedded and sliced into 3-μm sections for histology.

### Histology and Immunohistochemistry

Sections were deparaffinized in xylene, rehydrated in graded alcohol, and used for histology and immunostaining. Human tissue microarray was custom-made at Dr. Gaboury’s laboratory from the selected breast cancer cases, with each case represented in triplicate. Immunohistochemistry and immunofluorescence was performed with the antibodies listed in Table S3. In IHC experiments, detection was carried out using Streptavidin-HRP (1:1000 dilution, 554066, BD Pharmingen), or M.O.M. kit (Vector Laboratories), followed by Vectastain ABC (Vector Laboratories) according to the manufacturer’s instructions. For fluorescent image acquisition the exposure time in individual channels was determined by imaging software NIS-elements using filter sets: blue for DAPI/Hoechst (d460/50m); green for FITC, GFP, YFP, Alexa-488 (hq535/50m), red with narrow band excitation and red shifted emission for RFP, Alexa-568, Alexa-555 (hq620/60m) and far red cy5 filter for Alexa-647 (d680/30m). In some cases where the saturation of fluorescence signal was expected, exposure times were manually shortened. Pseudocolor images were created using Adobe Photoshop software. Bright-field images were captured using a Leica DMLB microscope, and images were processed with Magnavision or Adobe Photoshop software.

### Mathematical model

We have generated a schematic representation of the birth-death process in the lungs (Fig 3H). Using this scheme, we denote

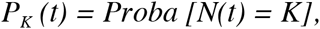
where *N(t)* is a number of marked cells at time *t* in the lungs, and *K* is an integer value. We define λ_K_ as the birth rate when there are K cells in the lungs and b_K_ as the death rate when there are K cells.

During a small interval of time *dt*, many scenarios can happen between *t* and *t* + *dt*, so that

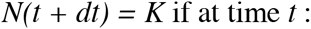

– Event 1: there were K − 1 cells and 1 is born; This event occurs with probability λ_*K*-*1*_*dt*.

– Event 2: there were K + 1 cells and 1 cell dies; This event occurs with probability b_*K*+*1*_*dt*

– Event 3: there were already K cells and 0 cells are born or died; This event occurs with probability 1-λ_*K*_*dt*-*b*_*K*_*dt*.

– Event 4: any other event different from events 1,2,3, as for instance the simultaneous birth or death of multiple cells. The probability of this event is *o*(*dt*), which is assumed to be extremely small.

Since birth can occur either from inside (by cell division in the ectopic breast cells in the lungs) or from outside (migration to the lung from the breast), one can decompose λ*_K_* as 
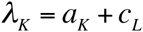
where

*a_K_* = birth rate of the cells inside the lungs when there are *K* cells in the lungs

*c_L_* = migration rate from the breast when there are *L* contacts in the breast

We summarize the previous birth and death possibilities in the following equation:

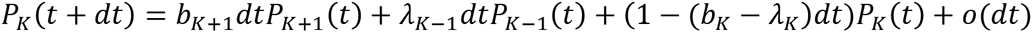

Now, we subtract *P_K_* (*t*), divide by *dt*, and take the limit when *dt* → 0; to obtain

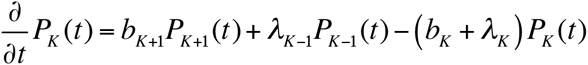

We assume that the birth and death rates in the lungs are the same for all the cells, that is, each cell in the lung is born at the same rate *a*, and dies at the same rate *b*, so we can write

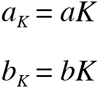

So that the dynamic of the cell’s growth may be written (for *K* ≥ 1)

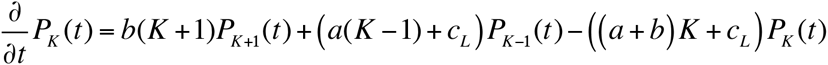

**Remark 1:** the boundary equation is when K = 0; where we have

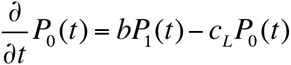

Justification: during a small interval of time *dt*, many scenarios can happen between *t* and *t* + *dt*, so *N(t + dt)* = *0* if at time *t*:

– Event 1: there was 1 cell and it died;

– Event 2: there were 0 cells and 0 birth;

– Event 3: any other event different from events 1,2.

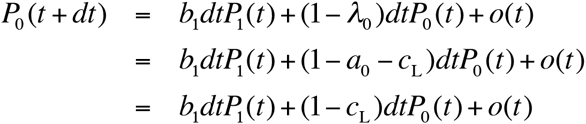
subtract *P_0_*(*t*), divide by *dt*, and take the limit when *dt*→0; to obtain the equation of 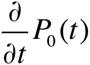

**Remark 2:** Since there is constant migration from the breast to the lungs, the cells in the lungs will only go extinct if the rate *c_L_* tends to 0:

**Remark 3:** The expected number of cells at time *t*, denoted *E[N(t)]*; is easier to compute since it satisfies

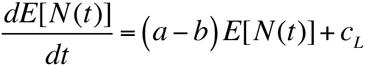

If we suppose that the initial number of cells in the lungs is *E[N(0)] = i*, then the expected number of cells in the lungs at time *t* is given by

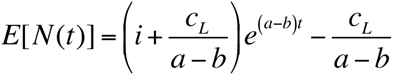

Note that the first term will tend to 0 when t increases, since a < b. As such after a few time steps the equation will tend to 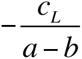. Thus with a constant in-flow of cells from the breast to the lungs the limit value will be some multiple of *c_L_* (see Fig 3I).

### Quantification of HA-stained sections

Tumor section were inspected microscopically and scored by the investigators as 100%, 90%, 75%, 50%, 25% positive, or as negative. One section was used for scoring the fully concordant and partially discordant cases, and three sections were used to score the fully discordant sections.

### Quantification of HA-lγH2A.X-double-stained sections

Early RCAS-csErbB2 lesion sections from three mice RCAS-ErbB2-injected and three control mice were inspected microscopically and composite fluorescent images were taken from 7 independent lesions and 5 uninjected ducts. 30 to 128 HA+, 12 to 33 HA-and 41 to 77 cells from unaffected ducts were evaluated per slide, and percent cells containing γH2A.X dots was calculated.

### QPCR of provirus

Genomic DNA was extracted from lung paraffin section as described (Hein and Doll 2005), and amplified with TaqMan^®^ qPCR oligos listed in Table S4. Relative load of provirus was determined as ΔCT against β-*actin* control.

### Statistical analyses

All of the data are presented as the mean ± standard errors of mean (SEM). All statistical analyses of mouse data have been performed with Prism 5 for MacOS X version 5.0b. Student’s *t* test or Mann-Whitney test were used to calculate the *p* values.

## Acknowledgements

We thank Myléne Cawthorn, Caroline Dube and Frederic Bourque for handling of the mouse colony; Jennifer Tran, Yildian Diaz-Rodriguez and Yasmina Hachem for genotyping of the mouse colony, Dominic Fillion for confocal microscopy, Marie Kmita, Jennifer Estall and Marina Glukhova (Institut Curie, France) for tissues from *mT/mG* mice, Philippe Chavrier (Institut Curie, France) for helping with animal housing, flow cytometry and microscopy charges, Christine Sedlik (Institut Curie, France) for 4T1 cell line and NSG mice, Lewis Chodosh (University of Pennsylvania School of Medicine, USA) for *TAN* and *MTB* mice. This work was supported by IRCM start-up funds and CIHR COP-126446 (to KP); PJA 20141201668 (to GDJ), and by NIH CA124820, CA202227, U54CA149196 (PI: Stephan Wong), DOD CDMRP BC133730, Sue & Lester Breast Center P50CA058183 & P50-CA186784, Dan L. Duncan Cancer Center P30CA125123 (to YL).

